# Serotonin and early life stress interact to shape brain architecture and anxious avoidant behavior – a TPH2 imaging genetics approach

**DOI:** 10.1101/685099

**Authors:** Congcong Liu, Lei Xu, Jialin Li, Feng Zhou, Xi Yang, Xiaoxiao Zheng, Meina Fu, Keshuang Li, Cornelia Sindermann, Christian Montag, Yina Ma, Dirk Scheele, Richard P. Ebstein, Shuxia Yao, Keith M. Kendrick, Benjamin Becker

## Abstract

**Background:** Early life stress has been associated with emotional dysregulations and altered architecture of limbic-prefrontal brain systems engaged in emotional processing. Serotonin regulates both, developmental and experience-dependent neuroplasticity in these circuits. Central serotonergic biosynthesis rates are regulated by Tryptophan hydroxylase 2 (*TPH2*), and transgenic animal models suggest that *TPH2*-gene associated differences in serotonergic signaling mediate the impact of aversive early life experiences on a phenotype characterized by anxious avoidance.

**Methods:** The present study employed an imaging genetics approach that capitalized on individual differences in a *TPH2* polymorphism (703G/T; rs4570625) to determine whether differences in serotonergic signaling modulate the effects of early life stress on brain structure and function and punishment sensitivity in humans (n = 252).

**Results:** Higher maltreatment exposure before the age of 16 was associated with increased gray matter volumes in a circuitry spanning thalamic-limbic-prefrontal regions and decreased intrinsic communication in limbic-prefrontal circuits selectively in TT carriers. In an independent replication sample, associations between higher early life stress and increased frontal volumes in TT carriers were confirmed. On the phenotype level, the genotype moderated the association between higher early life stress exposure and higher punishment sensitivity. In TT carriers, the association between higher early life stress exposure and punishment sensitivity was critically mediated by increased thalamic-limbic-prefrontal volumes.

**Conclusions:** The present findings suggest that early life stress shapes the neural organization of the limbic-prefrontal circuits in interaction with individual variations in the *TPH2* gene to promote a phenotype characterized by facilitated threat avoidance, thus promoting early adaptation to an adverse environment.

## Introduction

Early life stress (ELS) has been associated with strongly increased prevalence rates for psychiatric disorders and emotional dysregulations in adulthood. Converging evidence from animal and human studies suggest an association between ELS and lasting alterations in the structural and functional organization of the brain, particularly in limbic systems engaged in threat and anxiety processing and, including the amygdala-hippocampal complex, and frontal systems engaged in emotion regulation (Teicher, Samson, Anderson, & Ohashi, 2016).

On the behavioral level, considerable variations in the long-term sequelae of ELS have been reported. For instance, longitudinal data suggests that whereas the majority of individuals with severe ELS exhibits cognitive and emotional dysfunctions in adulthood, a significant minority remained unaffected (Sonuga-Barke et al., 2017), suggesting that innate factors such as genetic variations may temper the long-term trajectory of ELS. Initial studies that combined candidate gene approaches with neuroimaging reported that specific single-nucleotide polymorphisms (SNPs) of genes associated with neuroplasticity and stress reactivity (brain derived neurotrophic factor, oxytocin receptor) may confer differential susceptibility to the detrimental impact of aversive childhood experiences on the brain, i.e. differential reductions of limbic volumes following ELS exposure (Dannlowski et al., 2016; van Velzen et al., 2016).

Serotonin regulates both, neuromaturation during sensitive periods and learning-related neuroplasticity (Lesch & Waider, 2012) suggesting that innate differences in central serotonergic signaling may modulate the long-term effects of ELS. Central serotonin biosynthesis is regulated by the rate-limiting enzyme tryptophan hydroxylase (*TPH*), with the second isoform *TPH2* being exclusively expressed in central serotonin neurons (Zhang, Beaulieu, Sotnikova, Gainetdinov, & Caron, 2004).

Individual differences in the *TPH*2 encoding gene have been linked to variations in both, serotonin synthesis rates in limbic and prefrontal systems (Ottenhof, Sild, Levesque, Ruhe, & Booij, 2018; Furmark et al., 2016; Booij et al., 2012) and behavioral variations in emotional reactivity and regulation (Gutknecht et al., 2007; Laas et al., 2017; Reuter, Kuepper, & Hennig, 2007). Initial studies employing genetic imaging approaches furthermore reported that *TPH2* variants affect emotional reactivity in limbic regions and regulatory control in frontal regions (Canli, Congdon, Gutknecht, Constable, & Lesch, 2005; Canli, Congdon, Todd Constable, & Lesch, 2008; Kennedy et al., 2012; Reuter et al., 2008).

Transgenic animal models that capitalized on the selective effects of the *TPH2* isoform on central serotonin (e.g. in a mouse model with a mutated *TPH2* knock-in human polymorphism resulting in reduced brain 5-HT levels) suggest that congenital *TPH2*-associated serotonergic deficiency mediates the impact of early aversive experiences on anxious behavior in adulthood (Sachs et al., 2013), possibly due to serotonergic modulation of synaptic plasticity in limbic-prefrontal circuits that underpin emotional learning (Lesch & Waider, 2012). In humans *TPH2* SNP rs4570625 (−703 G/T) variations modulate the impact of ELS on emotional behavior, such that T-allele carriers exhibited increased threat attention in infanthood (Forssman et al., 2014) and elevated stress-reactivity in adulthood (Mandelli et al., 2012).

Despite accumulating evidence suggesting that *TPH2* genetics interact with early aversive experience to shape a phenotype characterized by anxious behavior, the brain systems that mediate this association have not been determined in humans.

To determine interactive effects of ELS and the *TPH2* SNP rs4570625 (−703 G/T) polymorphism on brain structure and function and their relationship to anxious-avoidant behavior in adulthood a sample of healthy subjects (n = 252) underwent *TPH2* rs4570625 genotyping and brain structural and functional magnetic resonance imaging (MRI). ELS exposure was retrospectively assessed by self-report, anxious behavior was assessed using a scale related to the Behavioral Inhibition System (BIS, Gray, 1987). The BIS appears particularly suitable given that it (1) specifically measures reactivity towards stimuli that are associated with a high probability of punishment and shapes anxious-avoidant behavior, and, (2) has been associated with the septo-hippocamal-amygdala system (Gray & McNaughton, 2000) and – in humans - prefrontal systems engaged in threat detection and emotion regulation (Fung, Qi, Hassabis, Daw, & Mobbs, 2019). Based on previous findings suggesting that (a) ELS is associated with brain structural and functional changes in limbic-frontal circuits and that (b) *TPH2* genetic variations are linked with serotonergic neurotransmission in these circuits (albeit with little direct evidence for the functional role of these SNPs (Ottenhof et al., 2018) and (c) brain serotonin levels modulate experience-dependent synaptic plasticity in limbic-frontal circuits, we hypothesized that, (1) the impact of ELS on the structural and functional organization of limbic-prefrontal circuits varies as a function of *TPH2* rs4570625 genetic differences, and (2) ELS-associated neuroplastic changes in this circuitry determine variations in anxious-avoidant behavior during early adulthood.

In line with previous studies on the effects of ELS on brain structure (e.g. Teicher et al., 2016) we furthermore expected a negative association between ELS and brain structure independent of genotype. Given the lack of previous studies on general effects of variations in *TPH2* on brain structure, no specific hypotheses in this respect were formulated. Based on recent meta-analytic evidence on the low robustness of interaction effects between candidate allele variants and ELS on complex behavioral traits (Border et al., 2019; Culverhouse, Saccone, & Bierut, 2018), we included an independent dataset to test the robustness of the findings observed in the discovery sample (see recommendations in Carter et al., 2017).

## Methods

### Participants

*N* = 252 healthy subjects (age range 18-29 years) served as discovery sample. In the context of problematic effect size determination in genetic imaging studies (Carter et al., 2017; Avinun, Nevo, Knodt, Elliott & Hariri, 2018) sample size was based on previous studies (Dannlowski et al., 2016; Van Velzen et al., 2016). In line with recent recommendations a replication sample was included (Carter et al., 2017). An independent sample from a study examining *TPH2* genotype and acute tryptophan depletion served as replication dataset (https://clinicaltrials.gov/ct2/show/NCT03549182, ID NCT03549182). The dataset included early life stress and brain structural assessments in healthy male *TPH2* rs4570625 homozygotes (TT, GG), data from n = 51 placebo-treated subjects served as replication dataset (details **Supplementary Information** and **Supplementary Table S1**). In both samples, participants with current or a history of medical, neurological/psychiatric disorders, regular use of psychotropic substances and MRI contraindications were excluded. The study was approved by the local ethics committee and in accordance with the latest revision of the Declaration of Helsinki. Written informed consent was obtained.

The discovery data represents part of the Chengdu Gene Brain Behavior Project that aims at determining behavioral and genetic associations in a large healthy cohort. The project incorporates studies focusing on the genetic and neural correlates of trait autism (Montag et al., 2017, Li et al., 2019) as well as determining interactions between early life stress exposure and *TPH2* genetics. The variables examined for the latter research aim have been prespecified in line with the study hypotheses. All variables, data exclusion and sample sizes are completely reported.

### Childhood experience, perceived stress and punishment sensitivity

Levels of ELS exposure were assessed using the Childhood Trauma Questionnaire (CTQ) (Bernstein et al., 2003) which consists of 25 retrospective self-report items spanning five types of aversive childhood experiences. In accordance with previous studies (e.g. Dannlowski et al., 2016), CTQ total score were used. Current stress (during the last months) was assessed using the Perceived Stress Scale (PSS, Cohen, Kamarck, & Mermelstein, 1983). The Sensitivity to Punishment scale (SPS, Sensitivity to Punishment and Sensitivity to Reward Questionnaire, Torrubia, Avila, Moltó, & Caseras, 2001) was administered to assess individual differences in behavioral inhibition. Individuals with high punishment sensitivity experience high levels of anxiety and avoid potential dangers (Gray, 1987). Cronbach’s α in the present sample demonstrated good internal consistency (0.847 CTQ; 0.834 PSS; 0.844 SPS). CTQ scores were non-normally distributed (*p* < 0.001, Shapiro-Wilk).

### Genotyping and MRI data

Genomic DNA was purified from buccal cells using a MagNA Pure96 robot (Roche Diagnostics, Mannheim), sample probes were designed by TIB MolBiol (Berlin, Germany). Genotyping was conducted by real-time polymerase chain reaction (RT-PCR) and subsequent high resolution high-resolution melting on a Cobas Z480 Light Cycler (Roche Diagnostics, Mannheim, Germany) according to the manufacturer’s instructions. MRI data acquisition was conducted on a 3.0 Tesla GE MR750 system (General Electric Medical System, Milwaukee, WI, USA) using standard acquisition parameters and procedures. Following initial quality assessments, the data was preprocessed using validated procedures in SPM12 (http://www.fil.ion.ucl.ac.uk/spm/), see **Supporting Information**.

### Analysis: genotype x ELS effects on brain structure and functional connectivity

Due to the non-normal distribution of CTQ scores, non-parametric permutation tests were employed to examine the effect of *TPH2* rs4570625genotype and ELS on brain structure and function (Permutation Analysis of Linear Models, PALM) (Winkler, Ridgway, Webster, Smith, & Nichols, 2014). Gray matter (GM) volumes and functional connectivity maps were entered into a single full factorial model respectively, with genotype group as between-subject factor (TT vs. TG vs. GG). CTQ scores were entered as covariate (dimensional model), including the crucial CTQ times group interaction term. In line with previous genetic imaging studies examining interactions between candidate genes and ELS on the brain (Dannlowski et al., 2013; Frodl et al., 2014; an Velzen et al., 2016), age, gender, education years, and for brain structure total intracranial volume, and for resting state mean framewise displacement were included as covariates. Based on the results of brain structural analysis we further explored whether regions showing *TPH2* genotype and ELS effects on brain structure additionally exhibit altered intrinsic functional interactions with other brain regions. To this end, seed-to-whole-brain voxel-wise connectivity maps were computed using standard protocols (Resting-State fMRI Data Analysis Toolkit; REST; http://www.restfmri.net, **Supporting Information**).

Statistical significance was determined using permutation-based inferences (10,000 permutations) and *p* _FWE_ < 0.05 using threshold-free cluster enhancement (TFCE) to control for multiple comparisons (within a gray matter mask, spm grey.nii > 0.3). In line with a previous study (Birn, Roeber, & Pollak, 2017), relative effects of current stress on the brain were determined by recomputing the analyses and substituting CTQ with PSS scores.

### Associations with anxious avoidant behavior

Associations between anxious avoidant behavior, ELS and brain structure or function (extracted gray matter and functional connectivity estimates from 5mm-radius spheres centered at the peak coordinates of interaction clusters) were next examined. The moderation effect of the *TPH2* rs4570625genotype on the association of CTQ with sensitivity to punishment was explored using a moderation analysis (PROCESS, Preacher & Hayes, 2004) estimating whether the interaction between the predictor (CTQ) and moderator (genotype) could predict the dependent variable (SPS scores). Next associations between ELS-associated brain structural or functional changes and the behavioral phenotype were explored using non-parametric correlation analyses (PALM, 10,000 permutations). Finally, a mediation analysis tested whether brain structural changes mediated effects of ELS on an anxious avoidant behavior (punishment sensitivity) within genotype groups (bootstrapping with 10,000 permutations, bias-corrected 95% confidence intervals (CIs) to test the significance of indirect effects, Preacher & Hayes, 2008).

### Replication procedure

Gray matter volumes from regions identified in the discovery sample were extracted from the MRI data of the replication sample and subjected as dependent variables to a partial correlation analyses with CTQ scores as independent variable (covariates: age, total intracranial volume).

### Data availability statement

The data that support the findings of the present study are openly available in the repository of the Open Science Framework at https://osf.io/DG8V3/, doi 10.17605/OSF.IO/DG8V3.

## Results

### Sample and genotyping

229 right-handed Han Chinese (age M=21.55 ±2.31, 115 females) were included in the final analysis (exclusion **Supplementary figure S1**). Genotyping for *TPH2* rs4570625 yielded *n* = 71 TT homozygotes, *n* = 107 GT heterozygotes, and *n* = 51 GG homozygotes. Allele frequencies did not deviate from those expected according to the Hardy-Weinberg Equilibrium (*χ*^2^_(1)_=0.735, *p*=0.391) and were in the expected distribution for Han Chinese populations (see https://www.ncbi.nlm.nih.gov/snp/rs4570625#frequency_tab). Previous studies on the *TPH2* rs4570625 were mostly conducted in Caucasian populations with a rare occurrence of the T-allele, in contrast the present distributions in Han Chinese allowed to separately examine GT and TT carriers. Genotype groups had comparable socio-demographics and CTQ scores (*p*s>0.36), however, PSS scores were significantly higher in TT compared to GT carriers (*F*_(2, 226)_=4.89, *p*=0.008, partial η2=0.041) Post hoc test revealed that PSS in TT homozygotes was significantly higher than in GT heterozygotes (*t*_(176)_=3.10, *p*=0.002, Cohen’s *d*=0.47) (**Table 1**).

**Table 1.** Sample characteristics (n = 229) stratified according to *TPH2* rs4570625

|  | TT (n = 71) | GT (n = 107) | GG (n = 51) | *p* Value | Total sample |
| --- | --- | --- | --- | --- | --- |
| Age, Years | 21.55 ±2.48 | 21.46 ±2.24 | 21.76 ±2.27 | 0.74<sup>a</sup> | 21.55 ±2.31 |
| Gender (M/F) | 32/39 | 56/51 | 26/25 | 0.63<sup>b</sup> | 114/115 |
| Education, Years | 15.55 ±2.09 | 15.69 ±1.84 | 15.78 ±1.84 | 0.79<sup>a</sup> | 15.67 ±1.91 |
| CTQ | 38.08 ±9.22 | 37.92 ±10.65 | 38.45 ±9.10 | 0.95<sup>a</sup> | 38.09 ±9.85 |
| PSS | 26.18 ±5.51 | 23.61 ±5.38 | 24.55 ±5.21 | 0.008<sup>a</sup> | 24.62 ±5.47 |
| SPS | 11.11 ±4.81 | 10.21 ±4.49 | 10.96 ±3.79 | 0.363<sup>a</sup> | 10.66 ±4.45 |

Mean and standard deviations ( $\pm SD$ ) are displayed. Abbreviations: CTQ, Childhood Trauma Questionnaire; F, female; M, male; PSS, Perceived Stress Scale; SPS, Sensitivity to Punishment scale. <sup>a</sup>*F* tests, <sup>b</sup> $\chi^2$ tests

### Interactive effects of TPH2 x ELS on brain structure

No significant main effects of ELS and genotype were found. Examining interactions of ELS x genotype revealed effects in a broad thalamic-limbic-frontal network, including a cluster spanning the bilateral thalamus, parahippocampal gyrus, hippocampus, and amygdala (k=2609, *p*FWE-TFCE<0.05), a cluster spanning the bilateral ventromedial frontal cortex (vmPFC) (k=1575, *p*FWE-TFCE<0.05), and additional clusters located in the left dorsolateral prefrontal cortex (DLPFC) (k=201, *p*FWE-TFCE<0.05), right (k= 335, *p*FWE-TFCE<0.05) as well as left (k=181, *p*FWE-TFCE<0.05) dorsal anterior cingulate cortex (dACC) (**Fig. 1**, details **Supplementary Table 2**). Post-hoc analyses by means of extracted parameter estimates from the significant interaction clusters revealed pronounced effects of ELS on gray matter volume in TT homozygotes, whereas associations in GT heterozygotes and GG homozygotes failed to reach significance. In TT homozygotes, higher stress exposure during childhood specifically associated with increased gray matter volumes in the bilateral thalamic-limbic cluster, left dACC and dlPFC (all *p*s<0.05 after Bonferroni correction, **Fig. 1**). Examination of correlation coefficients between the groups additionally revealed significant different associations between CTQ and volumes in these regions between TT and GT carriers (*p*corrected<0.01) (details **Supplementary Table 3**, effect sizes **Supplementary Table 4**). Findings remained stable after excluding the demographic covariates (**Supplementary Table 5**). No significant main and genotype interaction effects of current stress (PSS) were observed for brain structure.

**Fig. 1.**
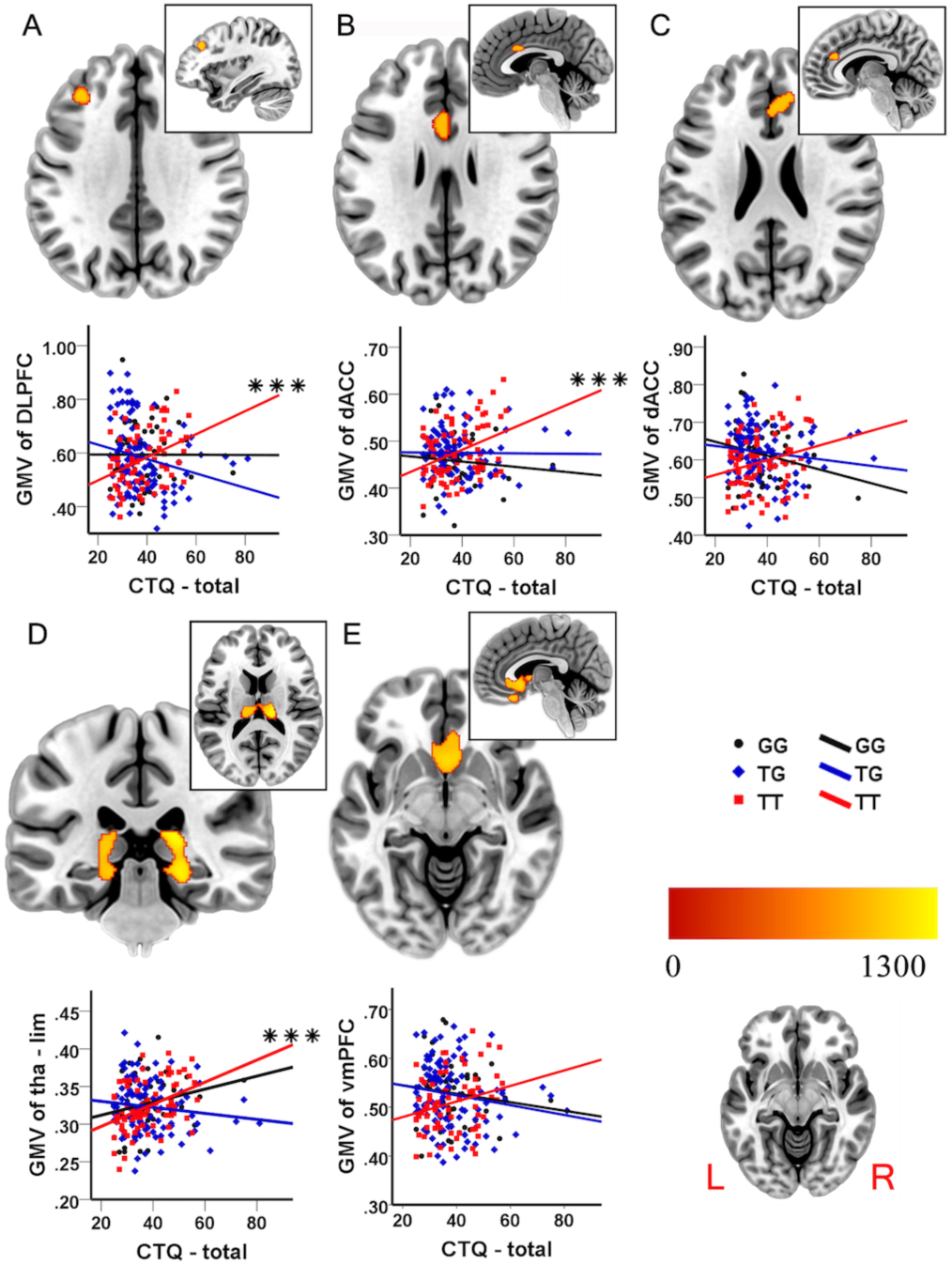
Interaction of *TPH2* rs4570625 and early life stress (ELS) on gray matter structure and relationship between gray matter volume and Childhood-Trauma-Questionnaire (CTQ) scores. Regions showing significant interaction effects between genotype and ELS (color bar = TFCE value, *p* _FWE with TFCE_ < 0.05) (2A-2E). CTQ, Childhood-Trauma-Questionnaire; dACC, dorsal anterior cingulate cortex; DLPFC, dorsolateral prefrontal cortex; ELS, early life stress; thalamic-limbic, parahippocampus/hippocampus/amygdala/thalamus; vmPFC, ventromedial frontal cortex; *** *p* < 0.001, Bonferroni-corrected

### Interactive effects of TPH2 x ELS on intrinsic network communication

ELS and *TPH2* rs4570625 interactively impacted functional connectivity strengths between the left dACC and left DLPFC (MNI_xyz_=[-6, 42, 27], k=24, *p*FWE-TFCE<0.05), with subsequent post-hoc tests demonstrating that the interaction was driven by pronounced effects of ELS in the TT homozygotes, whereas both GT and GG groups did not show significant associations (TT: *r*=-0.43, *p*_corrected_=0.0021, *d*=0.95; GT: *r*=0.11, *p*=0.27, *d*=0.22; GG: *r*=0.20, *p*=0.18, *d*=0.41; Fisher’s Z: *z*_TTvsGT_=3.61, *p*<0.001, *q*=0.57; *z*_TTvsGG_=3.47, p=0.001, *q*=0.66; *z*_GTvsGG_=0, *q*=0.09) (**Fig. 2**). Given previously reported genotype-dependent effects of ELS on amygdala activity (White et al., 2012) an amygdala-focused analysis (small-volume correction, SVC) was employed which revealed interactive effects on left dACC-left amygdala functional connectivity (MNI_xyz_ = [-21, 0, −12], k = 2, *p*_FWE_ < 0.05) (**Fig. 2**). The interaction was driven by a pronounced effect of ELS on left dACC-left amygdala connectivity in TT homozygotes, whereas GT and GG groups did not show significant associations (TT: *r*=-0.34, *p*_corrected_=0.032, *d*=0.0.72; GT: *r*=0.28, *p*_corrected_=0.055, *d*=0.58; GG: *r*=-0.14, *p*=0.35, *d*=0.28; Fisher’s Z: *z*_TTvsGT_=3.93, *p*<0.001, *q*=0.64; *z*_TTvsGG_=1.136, *p*=0.26, *q*=0.21; *z*_GTvsGG_=2.31, *p*=0.021, *q*=0.43) (**Fig. 2**). No significant interactive effects were observed for other seed regions. Findings remained stable after excluding the demographic covariates (**Supplementary Table 6**). No significant main and genotype interaction effects of current stress (PSS) were observed.

**Fig. 2.**
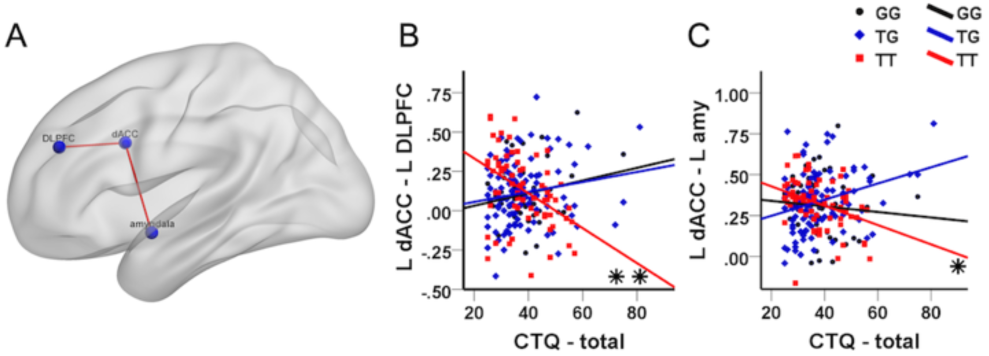
Interaction of *TPH2* rs4570625 and early life stress (ELS) on the functional connectivity strength of the left dorsal anterior cingulate cortex (dACC) (2A). Associations between extracted parameter estimates and Childhood-Trauma-Questionnaire (CTQ) total scores for dACC – left DLPFC (2B) left amygdala (2C). amy, amygdala; CTQ, Childhood-Trauma-Questionnaire; dACC, dorsal anterior cingulate cortex; DLPFC, dorsolateral prefrontal cortex; ELS, early life stress; * *p* < 0.05; ** *p* < 0.01, Bonferroni-corrected

### Associations with anxious avoidant behavior

The moderation analysis revealed a significant moderation effect (*R*^2^=0.05, *F*_(4, 224)_=3.12, *p*=0.016) reflecting that the interaction between CTQ and *TPH2* rs4570625 significantly predicts punishment sensitivity (*B*=0.094, *SE*=0.044, *t*_(224)_=2.15, *p*=0.03). Genotype thus significantly moderated the association between ELS and punishment sensitivity, with post hoc tests indicating that higher ELS selectively associated with higher punishment sensitivity in TT carriers (*t*_(224)_=2.59, *p*=0.01, (**Fig. 3**, visualization via Johnson-Neyman approach splitting CTQ scores into low, medium, high).

**Fig. 3.**
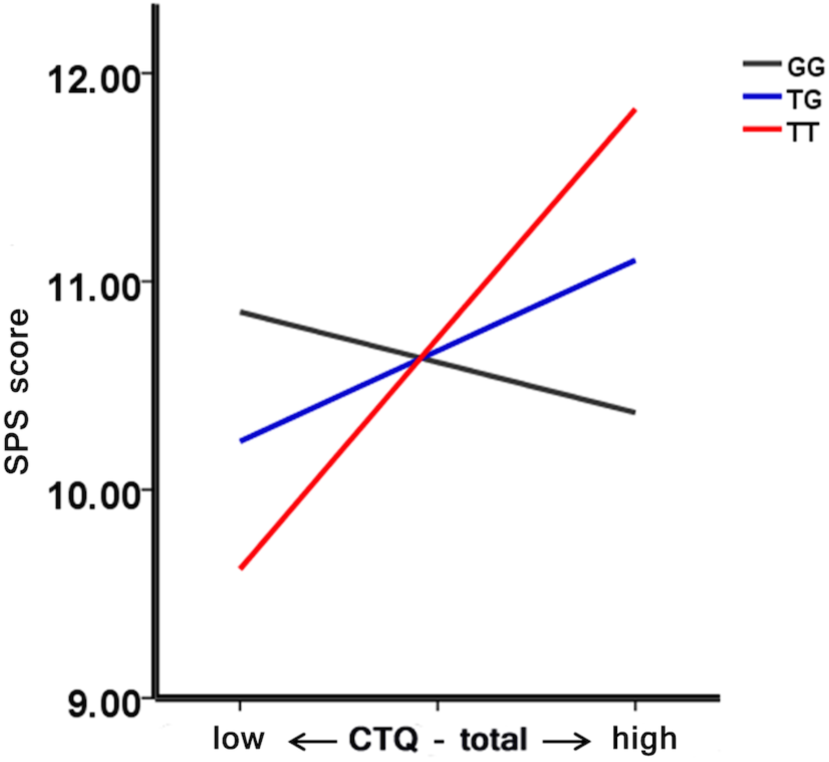
Moderation effect of genotype on the association between sensitivity to punishment scale (SPS) and Childhood Trauma Questionnaire (CTQ) scores. CTQ, Childhood Trauma Questionnaire; SPS, sensitivity to punishment scale;

Non-parametric correlation analysis revealed that specifically in TT homozygotes sensitivity to punishment was significantly positively associated with volumes of the thalamic-limbic region, right vmPFC and left DLPFC (vmPFC: *r*=0.359, *p*_corrected_=0.006, *d*=0.77; DLPFC: *r*=0.387, *p*_corrected_=0.008, *d*=0.84; thalamic-limbic: *r*=0.370, *p*_corrected_ =0.026, *d*=0.80) (**Fig. 4, Supplementary Table 8**). Associations between sensitivity to punishment and left dACC connectivity failed to reach significance and were not further examined.

**Fig. 4.**
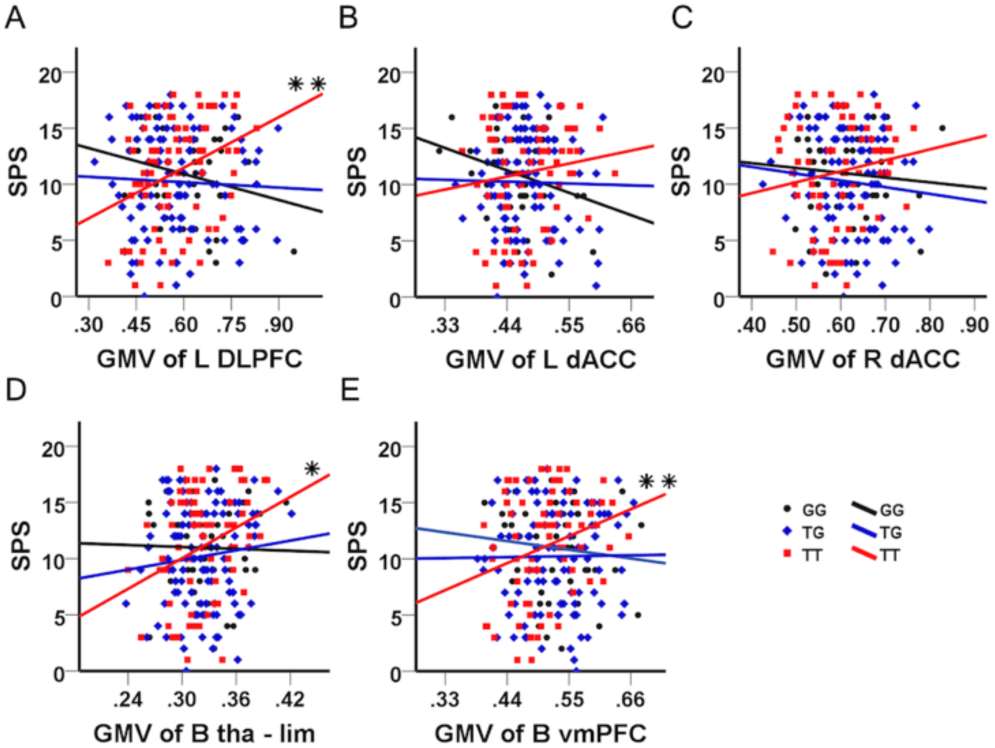
Relationship between gray matter volume and sensitivity to punishment scale (SPS) scores for TT homozygotes (red), TG heterozygotes (blue) and GG homozygotes (black) in the regions showing significant genotype x early life stress effects (4A-D). dACC, dorsal anterior cingulate cortex; DLPFC, dorsolateral prefrontal cortex; GMV, gray matter volume; SPS, sensitivity to punishment scale; vmPFC, ventromedial frontal cortex; thalamic-limbic, parahippocampus gyrus/ hippocampus/thalamus; L, left; R, right; B, bilateral; * *p* < 0.05; ** *p* < 0.01, Bonferroni-corrected

The mediation model reached significance in the TT group only, implying individual variations in sensitivity to punishment may be explained by the level of ELS exposure in the TT group. A significant overall indirect effect was observed (mediation by brain volume; indirect effect ratio 66.75%; effect size a×b=0.1096, 95%CI=[0.0247-0.2212]; *p*<0.05), suggesting that thalamic-limbic-prefrontal volumes mediated the influence of ELS on sensitivity to punishment in TT carriers (**Supplementary Table 9**).

### Replication sample

Results from the partial correlation analysis in the n = 51 male homozygotes revealed a similar pattern as in the discovery sample such that TT homozygotes exhibited a positive association between ELS and GMV (**Supplementary Table 7**). Specifically, in TT homozygotes, higher stress exposure during childhood associated with higher GMV in bilateral dACC and vmPFC. However, associations between higher ELS and increased volume in thalamic-limbic regions failed to reach significance in TT carriers, but exhibited a positive association in GG homozygotes. Thus, the positive associations between ELS and vmPFC and dACC volume in TT homozygotes appear to be robust, while associations in thalamic-limbic regions were less robust.

## Discussion

Previous animal models and initial findings in humans suggest that variations in serotonergic signaling associated with individual differences *TPH2* polymorphisms mediate the effects of early adverse experiences on emotional dysregulations (Mandelli et al., 2012, Sachs et al., 2013, Forssman et al., 2014). The current discovery sample found that *TPH2* rs4570625 genotype and ELS interact to shape the structural and functional architecture of thalamic-limbic-prefrontal circuits. Specifically, we observed that in TT homozygotes higher ELS exposure was associated with increased volumes of hippocampal-amygdala, thalamic and frontal regions and decreased functional connectivity between left dACC and left DLPFC, left amygdala, respectively. No associations were observed in the G-carrier groups. The replication dataset confirmed the robustness of the association between higher ELS and increased dACC and vmPFC volumes - at least for male subjects - whereas the observed associations in thalamic-limbic regions appeared to be less robust. Moreover, in the discovery sample *TPH2* genotype significantly moderated the impact of ELS on punishment sensitivity. In TT homozygotes higher levels of sensitivity to punishment were associated with both higher exposure to ELS and limbic and frontal brain volumes. An additional mediation analysis furthermore suggests that increased brain volumes in this circuitry critically mediate the impact of aversive childhood experiences on sensitivity to punishment solely in TT carriers.

In contrast to the present findings several previous studies that did not account for genetic differences, reported that higher ELS associated with decreased brain volumes in limbic and prefrontal regions (Teicher et al., 2016, Maier et al., 2020). An additional analysis that pooled the G-carriers of the present sample, revealed a negative association between ELS and brain volumes in the pooled G-carrier group partly replicating the previous findings (**Supplementary Table 10**). According to Cohen’s (1988) convention of effect sizes, the observed significant correlations in the pooled G-carrier group are small (between 0.1 and 0.3), while the associations in the TT group were medium to large (between 0.3 and 0.5). Animal models suggest that prolonged stress during sensitive developmental periods detrimentally impacts dendritic growth in limbic and prefrontal regions (Yang et al., 2015) which could partially explain lower volumes observed on the macro-anatomic level (Ivy et al., 2010). The few previous studies that examined whether polymorphisms modulate the impact of ELS on brain structure in humans observed that specific allelic variants coding brain-derived neurotrophic growth factor and oxytocinergic pathways pathway genes exhibit pronounced volumetric decreases following ELS (Dannlowski et al., 2016, van Velzen et al., 2016)

The observed opposite association between higher ELS and increased brain volumes observed in the *TPH2* rs4570625 TT homozygotes contrasts with these previous reports on a genetic susceptibility for pronounced ELS-associated brain structural deficits. Furthermore, some studies that combined the low-frequency *TPH2* rs4570625 T-carrier groups (TT and TG) initially revealed increased psychopathological risk (Gao et al., 2012), anxiety-associated traits (Gutknecht et al., 2007) and detrimental effects of ELS (Mandelli et al., 2012, Forssman et al., 2014). However, accumulating evidence indicates that TT homozygotes differ from both, TG and GG variants in terms of lower anxiety-related behavior and psychopathological risk (Ottenhof et al., 2018). While reduced gray matter volumes in limbic-prefrontal regions have been consistently reported in psychiatric disorders characterized by exaggerated anxiety and deficient emotion regulation (Bora, Fornito, Pantelis, & Yücel, 2012), associations between emotional functioning and limbic-prefrontal volumes in healthy subjects remain less clear, with elevated anxiety being associated with both volumetric increases and decreases in this circuitry (e.g. Günther et al., 2018). On the other hand, a growing number of studies consistently reported positive associations between aversive learning and volumetric indices in this circuit (e.g. Hartley, Fischl, & Phelps, 2011) and trauma exposure in adulthood has been associated with volumetric increases of the amygdala and concomitantly facilitated aversive learning (Cacciaglia et al., 2017). In the present study, both, childhood aversive experience and thalamic-limbic-prefrontal volumes were positively associated with higher punishment sensitivity in the TT homozygotes. According to the reinforcement sensitivity theory, individual differences in trait punishment sensitivity are neurally underpinned by the BIS, a brain-behavioral system that facilitates the formation of aversive motivation by orchestrating behavior in response to conditioned aversive events (Gray, 1987, Gray & McNaughton, 2000) and interacts with associative learning to facilitate aversive learning and future avoidance of threats (Reynolds, Askew, & Field, 2018). Increased punishment sensitivity on the phenotype level is considered to reflect an underlying hyperactive BIS system and has been consistently associated with better threat detection and facilitated aversive and avoidance learning (Avila & Parcet, 2000). The identified network exhibiting genotype x ELS exposure interaction effects highly overlaps with the neural circuits engaged in these functional domains, specifically the acquisition and extinction of conditioned threat responses (Milad & Quirk, 2012). Within this circuitry, the amygdala, hippocampus and dACC detect and encode threat probability of stimuli in the environment (e.g. Rodrigues, LeDoux, & Sapolsky, 2009) and are regulated by prefrontal-limbic cicruits that subserve implicit and explicit regulation of the innate or acquired threat response (Kalisch, 2009, Ramanathan, Jin, & Giustino, 2018). Serotonin plays an important role in these aversive learning mechanisms and underpins neuroplasticity in limbic-prefrontal circuits (Lesch & Waider, 2012). Mice with a knockin mutilated *TPH2* polymorphism (rs4570625) resulting in serotonin deficiency and exhibit a phenotype characterized by facilitated threat reactivity and threat acquisition that are mediated by neural morphology in limbic regions, particularly the hippocampal-amygdalar complex (Waider et al., 2019).

Although the neurobiological implications of the *TPH2* rs4570625 polymorphism are not fully understood, underscored by the T-variant being associated with increased as well as decreased central serotonergic transmission in limbic-prefrontal circuits (Zhang et al., 2004, Lin et al., 2007). The present findings might reflect that the *TPH2* polymorphism, perhaps underlying individual differences in 5HT neurotransmission, may mediate the impact of ELS via experience-dependent neuroplastic changes in limbic-prefrontal circuits engaged in aversive learning. This suggests the notion that this polymorphism interacts with aversive environmental factors to shape a neural and behavior phenotype with an enhanced capability to detect and avoid threat and thus facilitating survival in a malevolent environment (Teicher et al., 2016).

On the network level, ELS-associated volumetric increases observed in TT carriers were accompanied by decreased functional communication of the dACC with both, the DLPFC and the amygdala. Deficient intrinsic functional interaction within this circuit has been increasingly reported following ELS and has been associated with increased emotional dysregulations and threat reactivity on the behavioral level (Kaiser et al., 2018). Deficient recruitment of the dACC and functional communication within the dACC-DLPFC-amygdala circuitry have been associated with impaired voluntary emotion regulation (Zilverstand, Parvaz, & Goldstein, 2017, Zimmermann et al., 2017). In this context, the present findings could suggest that the increased regional-specific volumes of the aversive learning circuit in TT carriers come at the expense of less regulatory control within this circuitry, shifting the systems towards higher sensitivity for reacting to threatening stimuli in the environment rather than focusing on internal emotion regulation. These alterations may reflect experience-dependent adaptations that promote avoidance of further harm in a malevolent childhood environment however may render individuals at an increased (latent) psychopathological vulnerability in adulthood. Although individuals with psychiatric diagnosis were not enrolled in the present study, TT carriers reported slightly elevated levels of perceived current stress possibly reflecting elevated threat reactivity or deficient emotion regulation respectively. Alterations in these domains have been associated with aberrant amygdala-frontal functional connectivity that normalize with symptom-reduction in stress-related disorders (Spengler et al., 2017). Behavioral interventions and neuromodulatory strategies have been shown to facilitate emotion regulation via modulating amygdala-prefrontal connectivity (Zhao et al., 2019) and similar mechanims could be harrnessed to alleviate increased psychopathlogical vulnerability following ELS.

In the current study, genotype groups did not differ in the anxiety-related trait (sensitivity to punishment) which is not consistent with previous studies (Gutknecht et al., 2007; Laas et al., 2017; Reuter et al., 2007). For instance, Gutknecht et al. found that the T allele was a risk allele for cluster B and cluster C personality disorders and an anxiety-related phenotype (harm avoidance) in patients with personality disorders. On the other hand, Reuter et al. (2007) found lower harm avoidance in healthy TT homozygotes. The inconsistent findings may be related to population, ethnic and cultural diversity. In addition, the current results found that in TT carriers, there was a positive correlation between ELS and sensitivity to punishment, suggesting that the difference in early life experience may contribute to the diverse findings observed by investigators.

By leveraging on the genotyping of an apparently functional and common *TPH2* SNP on central serotonergic neurotransmission in combination with MRI and trait assessments in a representative sample of healthy subjects, the present study allowed us to determine the role of polymorphic gene differences that modulate serotonergic signaling and mediate the molecular adaptation to early life stress on the neural and phenotype level. Owing to the low prevalence of the T-allele in Caucasian populations, previous studies combined T-carriers (Markett et al., 2017), whereas the higher frequency of the T-variant in Asian populations uniquely enabled us to additionally examine characteristics of the TT-homozygote variant. Nevertheless, the current findings need to be tempered by the following limitations: (1) Early life stress was retrospectively assessed; however, the reported level of ELS did not differ between the genotype groups, (2) Different effects of ELS during different developmental periods have been reported, however the retrospective assessment does not allow further determination of genotype-dependent differences in the onset of stress exposure. (3) The replication sample did not include resting state data, female subjects or TG carriers and hence, the current findings require further replication.

## Conclusions

The present findings suggest that ELS shapes the neural organization of the thalamic-limbic-prefrontal circuits in interaction with individual genetic variations in a *TPH2* polymorphism. The identified network strongly overlaps with the circuitry engaged in aversive learning and avoidant behavior. Previous studies suggest that serotonin facilitates aversive learning via promoting neuroplasticity in thalamic-limbic-prefrontal circuits. Within this context the present findings suggest that genetic differences in serotonergic signaling associated with a specific *TPH2* polymorphism partially determine the impact of early life aversive experiences on the structural and functional architecture of the brain and shape a phenotype adapted to avoid threat in a malevolent environment.

## Acknowledgments

This work was supported by the National Key Research and Development Program of China (2018YFA0701400); National Natural Science Foundation of China (91632117, 31530032, 31700998); Open Research Fund of the State Key Laboratory of Cognitive Neuroscience and Learning, Beijing Normal University; Science, Innovation and Technology Department of the Sichuan Province (2018JY0001).

