## Supplementary information for "Serotonin and early life stress interact to shape brain architecture and anxious avoidant behavior – a TPH2 imaging genetics approach"

**Authors** Congcong Liu et al.

### Supplementary methods

*Participants: independent replication sample*

The validation sample included 51 healthy male participants aged 19-27 years (mean ± *SD* 22.02±2.48) who were recruited in a study examining interactive effects of acute tryptophan depletion and *TPH2* genetics (https://clinicaltrials.gov/ct2/show/NCT03549182, ID NCT03549182). To reduce variance in the primary outcomes of this study, only male participants were recruited. Exclusion criteria were similar to those in the discovery sample and included (1) current or a history of physical, neurological, or psychiatric disorders, (2) current or regular use of licit (nicotine, alcohol, medication) or illicit psychotropic substances, (3) weight > 85 kilograms, (4) MRI contraindications, (5) cardiovascular disorders including high blood pressure, (6) history of allergic reactions, (7) contraindications for acute tryptophan depletion. Demographics were similar to the discovery sample (see Supplemental Table 1).

*MRI data acquisition*

T1-weighted high-resolution anatomical images were acquired with a spoiled gradient echo pulse sequence, repetition time (TR) = 5.9 ms, echo time (TE) = minimum, flip angle = 9°, field of view (FOV) = 256 × 256 mm, acquisition matrix = 256 × 256, thickness = 1 mm, number of slice = 156. For the resting state, a total of 210 functional volumes were acquired using a T2*-

weighted Echo Planar Imaging (EPI) sequence (TR = 2000 ms, TE = 30 ms, FOV = 240 × 240 mm, flip angle = 90°, image matrix = 64 × 64, thickness/gap = 3.4/0.6mm, 39 axial slices with an interleaved ascending order). During the resting-state acquisition, participants were instructed to lie still and to fixate a white cross centered on a black background while not falling asleep. In post-MRI interviews, none of the subjects reported having fallen asleep during the scan. OptoActive MRI headphones (http://www.optoacoustics.com/) were used to reduce acoustic noise exposure during MRI acquisition (Roozen, Koevoets, & Den Hamer, 2008).

*Preprocessing of brain structural data*

T1-weighted anatomical images were initially visually inspected to verify absence of anatomical abnormalities and image quality, and next manually reoriented to the anterior commissure – posterior commissure (AC-PC). The structural images were subsequently preprocessed using SPM12 (Statistical Parametric Mapping, http://www.fil.ion.ucl.ac.uk/spm/). The brain volumes were first segmented into gray matter (GM), white matter (WM) and cerebrospinal fluid (CSF) using the new unified segmentation approach (Ashburner and Friston, 2005, Malone et al., 2015). Second, a group-specific template based on all participants was created using DARTEL algorithm (Ashburner, 2007). Next, the segmented GM and WM images were iteratively registered via the fast diffeomorphic registration DARTEL algorithm to warp the GM and WM partitions on to the template and subsequently non-linearly normalized to MNI space. Finally, gray matter volumes were spatially smoothed with a Gaussian 8 mm FWHM (full width at half maximum) kernel. Total intracranial volume (ICV) was estimated and used as a covariate on the second level.

*Preprocessing of resting state functional MRI data*

To examine whether the regional brain structural alterations were accompanied by functional communication differences between the nodes, resting state data was additionally acquired. Resting-state functional time-series were preprocessed using SPM12. For each subject, the first ten volumes were discarded to allow for T1 equilibration effect and allow active noise cancelling by the headphones. The remaining functional images were slice-time corrected and realigned to the first image to correct for head motion. The EPI images were then co-registered to the T1-weighted structural images, normalized to Montreal Neurological Institute (MNI) standard space using the segmentation parameters from the structural images and interpolated to 3 × 3 × 3 mm voxel size. Normalized images were finally spatially smoothed with a 6 mm FWHM. Next, 24 head movement parameters (i.e., 6 head motion parameters, 6 head motion parameters one time point before, and the 12 corresponding squared items) (Friston et al., 1996) along with mean signals from WM and CSF were removed from the data through linear regression. Finally, the influences of low-frequency drift and high-frequency noise were restricted by a band-pass filter (0.01–0.1 Hz). For the rsfMRI analyses, three subjects were excluded due to excessive head motion (exclusion criterion> 3 mm and/or 3 degrees, n = 2) or missed rsfMRI data (n = 1).

Functional connectivity analyses were performed using the Resting-State fMRI Data Analysis Toolkit (REST; http://www.restfmri.net). 5-mm spheres centered at the peak coordinates of significant clusters in the VBM analyses served as seed regions to create seed-to-whole brain intrinsic connectivity maps. To this end, the individual mean time series of each seed ROI was calculated and next correlation coefficients between the seed time series and other voxels were calculated to obtain *r* maps for each participant. Fisher *z* score transformations were employed to generate *z*-FC maps. Finally, *z*-FC maps were subjected to second-level random-effects group analyses. Consistent with the brain structural analyses, a single full factorial model was conducted using PALM with genotype group (TT vs GT vs GG) as between-subjects factor. Childhood Trauma Questionnaire (CTQ) scores were entered as covariate, including the interaction term of genotype and CTQ. Age, gender, education and mean framewise displacement (Van Dijk et al., 2012) were included as nuisance regressors. All covariates were mean-centered across participants.

**References supplementary materials**


Ashburner, J. (2007). A fast diffeomorphic image registration algorithm. *Neuroimage, 38*(1), 95-113. doi: https://doi.org/10.1016/j.neuroimage.2007.07.007

Ashburner, J., & Friston, K. J. (2005). Unified segmentation. *Neuroimage, 26*(3), 839-851. doi: 10.1016/j.neuroimage.2005.02.018

Friston, K. J., Williams, S., Howard, R., Frackowiak, R. S., & Turner, R. (1996). Movement-re lated effects in fMRI time-series. *Magn Reson Med, 35*(3), 346-355.

Malone, I. B., Leung, K. K., Clegg, S., Barnes, J., Whitwell, J. L., Ashburner, J., . . . Ridgway, G. R. (2015). Accurate automatic estimation of total intracranial volume: a nuisance variable with less nuisance. *Neuroimage, 104*, 366-372.

Roozen, N., Koevoets, A., & Den Hamer, A. (2008). Active vibration control of gradient coils to reduce acoustic noise of MRI systems. *IEEE/ASME Transactions on Mechatronics, 13*(3), 325-334. doi: DOI: 10.1109/TMECH.2008.924111

Van Dijk, K. R., Sabuncu, M. R., & Buckner, R. L. (2012). The influence of head motion on intrinsic functional connectivity MRI. *Neuroimage, 59*(1), 431-438. doi: 10.1016/j.neuroimage.2011.07.044


**Supplementary figure 1** Flow diagram displaying exclusion of participant and rationale for exclusion

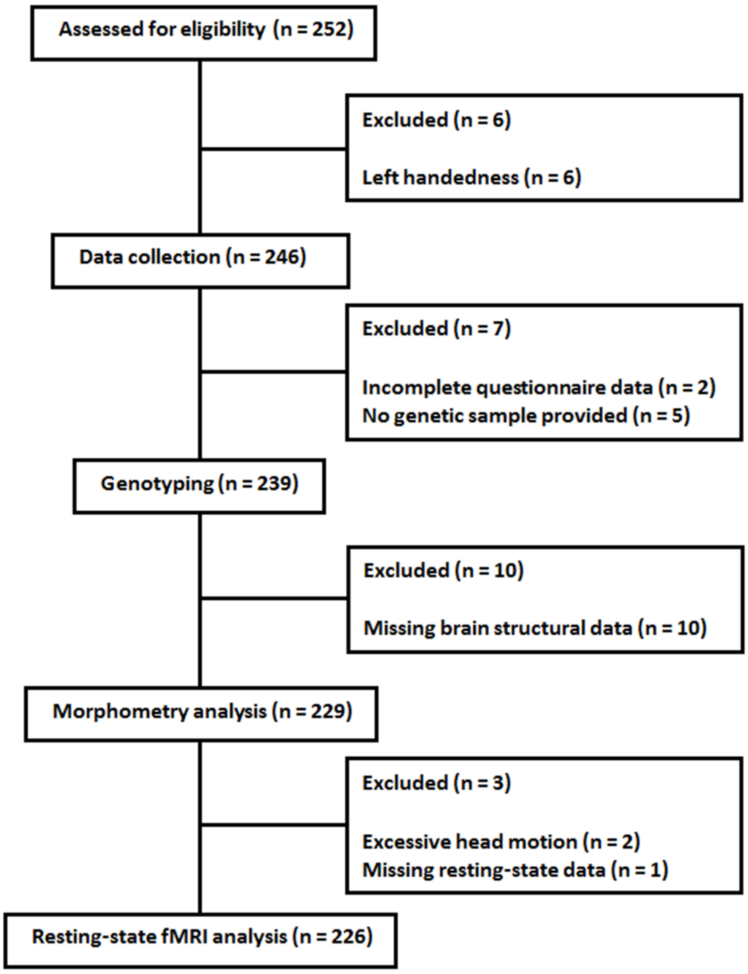


**Supplementary figure 2** distribution of the sensitivity to punishment scale (SPS) and Childhood Trauma Questionnaire (CTQ) scores.

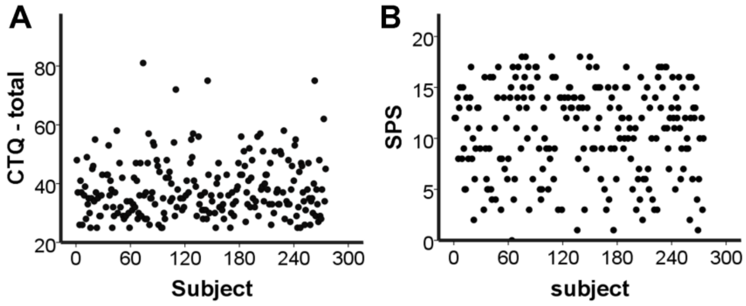


SPS, sensitivity to punishment scale; CTQ, Childhood Trauma Questionnaire.

**Supplementary table 1**

Sample characteristics in the discovery (n = 58 male subjects) and replication sample (n = 51)

| | Discovery study | | Replication study | | p value |
|---|---|---|---|---|---|
| | TT | GG | TT | GG | |
| Sample size | 32 | 26 | 27 | 24 | 0.52 |

| | | | | | |
|---|---|---|---|---|---|
| Age, Years | 21.55 ±2.48 | 21.76 ±2.27 | 22.07 ±2.29 | 21.96±2.72 | 0.35 |
| CTQ | 38.08 ±9.22 | 38.45 ±9.10 | 40.52 ±8.46 | 38.40 ±5.08 | 0.32 |

**Supplementary table 2**

Interaction between *TPH2* genotype and ELS on brain structure (whole-brain) in the discovery sample

| Brain Area | Peak located at MNI | | | TFCE | Cluster Size k | $p$ FWE with TFCE |
|---|---|---|---|---|---|---|
| L dACC | -1.5 | 12 | 28.5 | 932.25 | 181 | 0.045 |
| L DLPFC | -34.5 | 31.5 | 34.5 | 1008.65 | 201 | 0.034 |
| B thalamic -limbic | 19.5 | -25.5 | 12 | 1238.10 | 2609 | 0.015 |
| B vmPFC | 1.5 | 15 | -7.5 | 1068.18 | 1575 | 0.027 |
| R dACC | 6 | 33 | 24 | 956.25 | 335 | 0.042 |

Abbreviations: B, bilateral; dACC, dorsal anterior cingulate cortex; DLPFC, dorsolateral prefrontal cortex; GMV, gray matter volume; L, left; R, right; thalamic-limbic, parahippocampus gyrus/hippocampus/thalamus/amygdala; TFCE, threshold-free cluster enhancement; vmPFC, ventromedial frontal cortex.

**Supplementary table 3**

Correlation between GMV and CTQ and correlation differences between genotypes in the discovery sample

| Brain Area | MNI coordinates | | | Genotype[a] | | | *Z* value[a] | | |
|---|---|---|---|---|---|---|---|---|---|
| | | | | GG | TG | TT | GG *vs* TG | GG *vs* TT | TG *vs* TT |
| L dACC | -1.5 | 12 | 28.5 | 0.00 | -0.03 | 0.47*** | 0.19 | 2.71 | 3.48** |
| L DLPFC | -34.5 | 31.5 | 34.5 | 0.08 | -0.27 | 0.49*** | 2.08 | 2.38 | 5.22*** |
| B thalamic -limbic | 19.5 | -25.5 | 12 | 0.37 | -0.15 | 0.47*** | 3.11* | 0.65 | 4.26*** |
| B vmPFC | 1.5 | 15 | -7.5 | -0.03 | -0.25 | 0.35 | 1.27 | 2.07 | 3.93** |
| R dACC | 6 | 33 | 24 | -0.12 | -0.18 | 0.35 | 0.33 | 2.59 | 3.49** |

Mean gray matter volumes were extracted from a 5-mm sphere centered at the peak coordinates of the respective clusters identified in the VBM analyses. Results were corrected for multiple comparisons using Bonferroni correction [(a)].

Abbreviations: CTQ, Childhood Trauma Questionnaire; dACC, dorsal anterior cingulate cortex; DLPFC, dorsolateral prefrontal cortex; GMV, gray matter volume; L, left; R, right; thalamic-limbic, parahippocampus gyrus/hippocampus/thalamus/amygdala; vmPFC, ventromedial frontal cortex . * $p < 0.05$; ** $p < 0.01$; *** $p < 0.001$.

**Supplementary table 4**

Effect sizes of correlation between GMV and CTQ and correlation differences between genotypes

| Brain Area | MNI coordinates | | | Cohen's *d* | | | Cohen's *q* | | |
|---|---|---|---|---|---|---|---|---|---|
| | | | | GG | TG | TT | GG *vs* TG | GG *vs* TT | TG *vs* TT |
| L dACC | -1.5 | 12 | 28.5 | 0.00 | 0.06 | 1.06** | 0.03 | 0.51** | 0.54** |
| L DLPFC | -34.5 | 31.5 | 34.5 | 0.16 | 0.56* | 1.12 ** | 0.36* | 0.49* | 0.81** |
| B thalamic -limbic | 19.5 | -25.5 | 12 | 0.80* | 0.30 | 1.06** | 0.54* | 0.12 | 0.66** |
| B vmPFC | 1.5 | 15 | -7.5 | 0.06 | 0.51* | 0.75* | 0.23 | 0.40* | 0.62** |
| R dACC | 6 | 33 | 24 | 0.24 | 0.37 | 0.75* | 0.06 | 0.49* | 0.55** |

To explore the strengths of the associations and correlation differences effects sizes in terms of Cohen's d and q were computed.

Abbreviations: dACC, dorsal anterior cingulate cortex; vmPFC, ventromedial frontal cortex; DLPFC, dorsolateral prefrontal cortex; thalamic-limbic, parahippocampal gyrus/ hippocampus/thalamus /amygdala; L, left; R, right; GMV, gray matter volume; CTQ, Childhood Trauma Questionnaire. * medium effect; ** large effect size

**Supplementary table 5**

Correlation between GMV and CTQ with ICV as covariate and correlation differences between genotypes

| Brain Area | MNI coordinates | | | Genotype[a] | | | *Z* value[a] | | |
|---|---|---|---|---|---|---|---|---|---|
| | | | | GG | TG | TT | GG *vs* TG | GG *vs* TT | TG *vs* TT |
| L dACC | -1.5 | 12 | 28.5 | -0.06 | -0.03 | 0.45*** | 0.17 | 2.89 | 3.3* |
| L DLPFC | -34.5 | 31.5 | 34.5 | 0.12 | -0.28 | 0.49*** | 2.34 | 2.2 | 5.28*** |
| B thalamic -limbic | 19.5 | -25.5 | 12 | 0.37 | -0.14 | 0.46*** | 3.03* | 0.58 | 4.09*** |
| B vmPFC | 1.5 | 15 | -7.5 | -0.003 | -0.23 | 0.33 | 1.32 | 1.83 | 3.7** |
| R dACC | 6 | 33 | 24 | -0.13 | -0.18 | 0.33 | 0.29 | 2..51 | 3.37* |

Mean gray matter volumes were extracted from a 5-mm sphere centered at the peak coordinates of the respective clusters identified in the VBM analyses. Results were corrected for multiple comparisons using Bonferroni correction [(a)]. The primary findings remained robust after excluding gender, age and education as covariates.

Abbreviations: CTQ, Childhood Trauma Questionnaire; dACC, dorsal anterior cingulate cortex; DLPFC, dorsolateral prefrontal cortex; GMV, gray matter volume; ICV, intracranial volume; L, left; R, right; thalamic-limbic, parahippocampal gyrus/ hippocampus/thalamus/amygdala; vmPFC, ventromedial frontal cortex;. * $p < 0.05$; ** $p < 0.01$; *** $p < 0.001$.

**Supplementary table 6**

Correlation between functional connectivity values and CTQ with FD as covariates and correlation differences between genotypes

| Brain Area | MNI coordinates | | | Genotype[a] | | | Z value[a] | | |
|---|---|---|---|---|---|---|---|---|---|
| | | | | GG | TG | TT | GG | GG | TG |
| | | | | 51 | 106 | 69 | *vs* TG | *vs* TT | *vs* TT |
| L dACC - amygdala | -21 | 0 | -12 | -0.15 | 0.21* | -0.35* | 2.08 | 1.13 | 3.67*** |
| L dACC - L DLPFC | -6 | 42 | 27 | 0.24 | 0.14 | -0.43*** | 0.59 | 3.71** | 3.81*** |

Mean functional connectivity values were extracted from a 5-mm sphere centered at the peak coordinates of the respective clusters identified in the functional connectivity analyses. Results were corrected for multiple comparisons using Bonferroni correction [(a)]. The primary findings remained robust after excluding gender, age and education as covariates.
Abbreviations: CTQ, Childhood Trauma Questionnaire; dACC, dorsal anterior cingulate cortex; DLPFC, dorsolateral prefrontal cortex; FD, framewise displacement; L, left; R, right. * $p < 0.05$; ** $p < 0.01$; *** $p < 0.001$.

**Supplementary Table 7**

Correlation between GMV and CTQ in the discovery (n = 58 male subjects) and replication sample

| Brain Area | discovery sample | | replication sample | |
|---|---|---|---|---|
| | GG(n = 26) | TT (n = 32) | GG (n = 24) | TT (n = 27) |
| L dACC | -0.102 | 0.653*** | -0.178 | 0.42* |
| L DLPFC | 0.144 | 0.614*** | 0.372 | 0.246 |
| B tha-lim | 0.021 | 0.419* | 0.523* | 0.033 |
| B vmPFC | -0.141 | 0.499** | 0.186 | 0.454* |
| R dACC | -0.201 | 0.63*** | 0.084 | 0.552** |

Mean gray matter volumes were extracted from both samples using a 5-mm sphere centered at the peak coordinates of the respective clusters identified in the VBM analyses. Abbreviations: CTQ, Childhood Trauma Questionnaire; dACC, dorsal anterior cingulate cortex; DLPFC, dorsolateral prefrontal cortex; GMV, gray matter volume; L, left; R, right; thalamic-limbic, parahippocampus gyrus/hippocampus/thalamus/amygdala; vmPFC, ventromedial frontal cortex;. * $p < 0.05$; ** $p < 0.01$; *** $p < 0.001$.

**Supplementary Table 8**

Associations between gray matter volume and sensitivity to punishment in the discovery sample

| Brain Area | MNI coordinates | | | Genotype[a] | | | Z value[b] | | |
|---|---|---|---|---|---|---|---|---|---|
| | | | | GG | TG | TT | GG *vs* TG | GG *vs* TT | TG *vs* TT |
| L dACC | -1.5 | 12 | 28.5 | -0.14 | 0.02 | 0.20 | 0.90 | 1.83 | 1.21 |
| L DLPFC | -34.5 | 31.5 | 34.5 | -0.12 | -0.02 | 0.39** | 0.62 | 2.83** | 2.72** |
| B thalamic -limbic | 19.5 | -25.5 | 12 | 0.07 | 0.15 | 0.37* | 0.46 | 1.68 | 1.51 |
| B vmPFC | 1.5 | 15 | -7.5 | 0.07 | 0.06 | 0.36** | 0.02 | 1.65 | 2.01* |
| R dACC | 6 | 33 | 24 | 0.06 | -0.05 | 0.24 | 0.65 | 0.96 | 1.89 |

Gray matter volumes extracted from regions demonstrating significant environment x genetic interactions and associations with punishment sensitivity. Positive associations were specifically observed in the TT homozygotes.
Abbreviations: B, bilateral; GMV, grey matter volume; dACC, dorsal anterior cingulate cortex; DLPFC, dorsolateral prefrontal cortex; L, left; R, right; thalamic-limbic, parahippocampus gyrus/ hippocampus/thalamus/amygdala; SPS, sensitivity to punishment scale; vmPFC, ventromedial frontal cortex;.
* $p < 0.05$; ** $p < 0.01$; *** $p < 0.001$; [a]multiple comparisons corrected with Bonferroni correction; [b]no Bonferroni-correction applied

**Supplementary table 9**

Detailed results from the mediation analysis in the discovery sample

| Group | | Coefficient | *SE* | *p* | Bootstrap 95% CI | Effect ratio |
|---|---|---|---|---|---|---|
| TT | Model | $R^2$=0.304; F[7, 63]=3.932, *p*=0.001 | | | | |
| | Total effect | 0.164 | 0.059 | 0.007 | (0.046, 0.282)* | 100% |
| | Direct effect | 0.055 | 0.067 | 0.415 | (-0.078, 0.188) | 33.25% |
| | indirect effect(via mediators) | 0.110 | 0.048 | | (0.027, 0.220)* | 66.75% |
| | L dACC | -0.033 | 0.036 | | (-0.122, 0.024) | -19.78% |
| | L DLPFC | 0.049 | 0.029 | | (0.002, 0.118)* | 30.04% |
| | B thalamic-limbic | 0.049 | 0.028 | | (0.005, 0.117)* | 30.05% |
| | R vmPFC | 0.04 | 0.031 | | (-0.006, 0.116) | 24.36% |
| | R dACC | 0.003 | 0.027 | | (-0.055, 0.055) | 2.08% |
| TG | Model | $R^2$=0.055;F[7, 99]=0.821, *p*=0.572 | | | | |
| | Total effect | 0.001 | 0.041 | 0.989 | (-0.081, 0.082) | 100% |
| | Direct effect | 0.008 | 0.043 | 0.849 | (-0.078, 0.095) | 1454.34% |
| | indirect effect(via mediators) | -0.008 | 0.018 | | (-0.045, 0.030) | -1354.34% |
| | L dACC | -0.000 | 0.005 | | (-0.017, 0.008) | -52.01% |
| | L DLPFC | -0.003 | 0.017 | | (-0.043, 0.027) | 579.94% |
| | B thalamic-limbic | -0.011 | 0.010 | | (-0.042, 0.002) | 1949.37% |
| | R vmPFC | -0.008 | 0.014 | | (-0.046, 0.012) | 1432.01% |
| | R dACC | 0.015 | 0.015 | | (-0.003, 0.061) | 2658.99% |
| GG | Model | $R^2$=0.148;F[7, 43]=1.063, *p*=0.403 | | | | |
| | Total effect | -0.018 | 0.059 | 0.765 | (-0.136, 0.101) | 100% |
| | Direct effect | -0.027 | 0.066 | 0.684 | (-0.161, 0.107) | 153.19% |
| | indirect effect(via mediators) | 0.009 | 0.041 | | (-0.075, 0.090) | -53.19% |
| | L dACC | 0.002 | 0.016 | | (-0.021, 0.052) | -10.50% |
| | L DLPFC | -0.006 | 0.013 | | (-0.062, 0.008) | 32.92% |
| | B thalamic-limbic | 0.021 | 0.027 | | (-0.029, 0.080) | -116.86% |
| | R vmPFC | -0.000 | 0.014 | | (-0.035, 0.025) | 0.58% |
| | R dACC | -0.007 | 0.021 | | (-0.077, 0.017) | 40.67% |

Detailed results from the mediation analyses testing the mediation effect of brain volume on the association between earl aversive experiences and trait sensitivity to punishment

* coefficient greater than zero at $p < 0.05$

Abbreviations: B, bilateral; dACC, dorsal anterior cingulate cortex; DLPFC, dorsolateral prefrontal cortex; L, left; R, right; thalamic-limbic, parahippocampus gyrus/ hippocampus/thalamus/amygdala; vmPFC, ventromedial frontal cortex.

**Supplementary table 10**

Associations between brain volume and early life stress in the combined G-carrier group in the discovery sample

| Brain Area | MNI coordinates | | | Genotype | |
|---|---|---|---|---|---|
| | | | | G-carrier | TT |
| L dACC | -1.5 | 12 | 28.5 | -0.02 | 0.47*** |
| L DLPFC | -34.5 | 31.5 | 34.5 | -0.16* | 0.49*** |
| B thalamic -limbic | 19.5 | -25.5 | 12 | 0.00 | 0.47*** |
| B vmPFC | 1.5 | 15 | -7.5 | -0.18* | 0.35** |
| R dACC | 6 | 33 | 24 | -0.17* | 0.35** |

Mean grey matter volumes were extracted from a 5-mm spheres centered at the peak coordinates of the respective clusters identified in the VBM analyses.

Abbreviations: CTQ, Childhood Trauma Questionnaire; dACC, dorsal anterior cingulate cortex; DLPFC, dorsolateral prefrontal cortex; GMV, gray matter volume;L, left; R, right; thalamic-limbic, parahippocampus gyrus/ hippocampus/thalamus/amygdala; ; vmPFC, ventromedial frontal cortex.

* $p < 0.05$; ** $p < 0.01$; *** $p < 0.001$.
